# Network analysis links genome-wide phenotypic and transcriptional stress responses in a bacterial pathogen with a large pan-genome

**DOI:** 10.1101/071704

**Authors:** Paul A. Jensen, Zeyu Zhu, Tim van Opijnen

**Affiliations:** Biology Department, Boston College, Chestnut Hill, MA, USA; Current address: Department of Bioengineering and Carl R. Woese Institute for Genomic Biology, University of Illinois at Urbana-Champaign, Urbana, IL, USA

**Author notes:** Equal contribution (Paul A. Jensen, Zeyu Zhu). Corresponding author, E-mail Addresses: PAJ, ZZ, TvO (corresponding).

**Keywords:** Tn-Seq 21, RNA-Seq, stress response, Streptococcus, systems biology, data integration, metabolic modeling

## Abstract

**Background:** Bacteria modulate subcellular processes to handle stressful environments. Genome-wide profiling of gene expression (RNA-Seq) and fitness (Tn-Seq) allows two views of the same genetic network underlying these responses. However, it remains unclear how they combine, enabling a bacterium to overcome a perturbation.

**Results:** Here we generate RNA-Seq and Tn-Seq profiles in three strains of *S. pneumoniae* in response to stress defined by different levels of nutrient depletion. These profiles show that genes that change their expression and/or become phenotypically important come from a diverse set of functional categories, and genes that are phenotypically important tend to be highly expressed. Surprisingly, we find that expression and fitness changes rarely occur on the same gene, which we confirmed by over 140 validation experiments. To rationalize these unexpected results we built the first genome-scale metabolic model of *S. pneumoniae* showing that differential expression and phenotypic importance actually correlate between nearest neighbors, although they are distinctly partitioned into small subnetworks. Moreover, a meta-analysis of 234 *S. pneumoniae* gene expression studies reveals that essential genes and phenotypically important subnetworks rarely change expression, indicating that they are shielded from transcriptional fluctuations and that a clear distinction exists between transcriptional and phenotypic response networks.

**Conclusions:** We present a genome-wide computational/experimental approach that contextualizes changes that occur on transcriptomic and phenomic levels in response to stress. Importantly, this highlights the need to connect disparate response networks, for instance in antibiotic target identification, where preferred targets are phenotypically important genes that would be overlooked by transcriptomic analyses alone.

## INTRODUCTION

How organisms handle and overcome stress in their environment is a central question in biology, with applications in fields ranging from bioengineering to drug(-target) discovery. With the advent of genome-wide approaches, it has become clear that even relatively simple perturbations, such as a change in extracellular pH or the presence of an antibiotic, require complex physiologic responses affecting multiple subcellular processes [1-7]. A complete cellular stress response can be separated into at least two organization levels. The *transcriptional response* describes the change in gene expression following stress, while the *phenotypic response* describes how the importance of each gene (fitness) changes due to stress. Although the transcriptional and phenotypic stress responses are inter-dependent, they are measured experimentally by two disparate technologies. Transcriptomic profiling, as measured by cDNA microarrays or RNA-Seq, has been particularly popular in deciphering the complex bacteria-environment interaction to identify genes that change in their transcript abundance upon various environmental perturbations, such as exposure to antibiotics [8,9], interaction with host niches [10], and disruption of iron homeostasis [11]. Although such profiles provide a detailed picture of the transcriptional landscape, it remains to be determined whether transcript abundance alone is predictive of the phenotypic importance of a gene [12,13]. Alternatively, genome-wide mutant fitness profiling approaches, such as transposon insertion sequencing (Tn-Seq), have been developed to directly link genotypes to phenotypes and thereby measure the phenotypic stress response on a genome-wide scale [14-18]. This means that Tn-Seq determines the phenotypic importance of each gene in the genome in a specific environment (referred to as “gene fitness” in this paper) by measuring the effect each gene knockout in the genome has on fitness. For example, the lower the fitness, the more important a gene is for maintaining survival under a specific (e.g. stressful) condition [2,14,18]. Although both transcriptomic and phenotypic fitness profiling interrogate the same gene network, little is known about how these two data types correlate, i.e. are differentially expressed genes also phenotypically important during stress? Indeed, it is generally assumed that differentially expressed genes also represent phenotypically important genes. However, this assumption has not been thoroughly investigated, raising the possibility that if transcriptional and phenotypic stress responses are not correlated, then measuring either one alone offers an incomplete and incorrect picture of the cellular response. Here we determine whether a correlation between transcription and phenotype exists and whether these stress responses can be combined to obtain a complete physiologic response.

As our model system we employ the human pathogenic bacterium *Streptococcus pneumoniae*, a major respiratory pathogen and source of morbidity and mortality. *S. pneumoniae* colonizes the nasopharynx asymptomatically, but by disseminating to other tissues it can trigger disease, including pneumonia, meningitis, sepsis, and otitis media, which results in ~1 million deaths annually among children <5 years of age and ~0.5 million among groups including the immunocompromised and the elderly (>65 years) [19-21]. One possible challenge in exploring the physiologic response in a species such as *S. pneumoniae* is the genomic variation among strains. The increasing availability of fully sequenced genomes for this and other species has demonstrated a distinction between the species’ core genome (the set of genes shared by all strains) and its pan-genome (the species’ global genetic repertoire) [22-25]. Because no gene or pathway functions in a vacuum, rather they are connected by complex genetic networks [26,27], the presence or absence of genes among different pneumococcal strains suggests that each strain’s genomic network may be differently wired, which could potentially make phenotypes strain-dependent. Indeed, we have recently shown that strain-specific phenotypic stress responses in *S. pneumoniae,* for instance in response to antibiotics, may be common [1]. Thus, when working with a bacterium with a large pan-genome, an ideal approach should obtain a species-wide, generalizable view of how the bacterium overcomes a stressful environment.

In this study, we investigate whether differentially expressed genes are also phenotypically important during stress in *S. pneumoniae.* We generate an unbiased, high-quality, and extensively validated profile of the bacterial stress response by employing two genome-wide approaches, RNA-Seq and Tn-Seq, to measure both the transcriptional and phenotypic stress response for three pneumococcal strains under three different levels of nutrient depletion. Surprisingly, there is little correlation between differential expression and phenotypic importance across the entire genome. To contextualize the transcriptional and phenotypic profiles, we built and curated the first genome-scale metabolic model in the genus *Streptococcus.* By integrating all our data into this model we show that the disparate genes with expression and fitness changes are actually closely connected in the genomic network. However, phenotypically important and essential genes seem to be transcriptionally shielded from large fluctuations in expression and therefore organizationally separated from transcriptionally plastic, but phenotypically unimportant, genes. Importantly, we provide a detailed roadmap to develop similar systems-level approaches in other microorganisms. Our approach facilitates profiling and reconciling transcriptomic and mutant fitness datasets and enables mapping of an organism’s full physiologic response to an environmental disturbance. Moreover, this study provides a clear rationale that emphasizes the importance of targeting phenotypically important genes rather than differentially expressed genes in applications such as in drug target discovery.

## RESULTS AND DISCUSSION

### Designing a robust nutrient depletion assay for *S. pneumoniae*

To avoid bias in the bacterial response to nutrient depletion that might result from genomic variation in a particular strain, we selected three phylogenetically distant strains to represent *S. pneumoniae*: TIGR4 (T4), Taiwan-19F (19F) and D39 (**Additional File 1**). Both T4 and 19F can cause invasive pneumococcal disease (IPD): T4 is a serotype 4 strain that was originally isolated from a Norwegian patient with IPD [28,29]; 19F is a multi-drug resistant strain isolated from a patient with IPD in Taiwan [30,31] (Table 1). D39 is a historically important and commonly used serotype 2 strain that was originally isolated from a patient about 90 years ago [32] (Table 1). Concerning their genomic content, the three strains share 1647 genes, while T4 has 217, 19F has 140 and D39 has 93 strain-specific genes (**Additional Files 2 and 3**). To simulate systemic nutrient depletion, we designed three increasingly restrictive media to cultivate *S. pneumoniae*. The first, semi-defined minimal media (SDMM), is a relatively rich media we have used previously [1,2] that contains a single carbon source (glucose), yeast extract, casein hydrolysate (digested amino acids), salts, trace metals, and vitamins (Table 2). The second, a chemically defined media (CDM), is based on a previously described recipe [33], however the original composition did not allow each strain to grow robustly. By adjusting the recipe, mainly by taking SDMM as the basis and replacing the yeast extract and casein hydrolysate by an equimolar mixture of the 20 amino acids, comparable growth rates to SDMM are achieved. CDM is thus completely defined, is less nutrient rich then SDMM, but still contains several non-essential components. By iteratively removing components of CDM, we created a minimal CDM, or MCDM, that still enables each strain to grow. In contrast to SDMM and CDM, each component in MCDM is essential for growth in at least one of the three strains, and removing any component of MCDM triggers severe growth defects in at least one strain. Nutrient availability therefore decreases from SDMM to CDM, and further to MCDM (**Additional File 4**), which is illustrated by a decrease in the growth rate of *S. pneumoniae* (Table 1, **Additional File 5**). By using multiple strains and three media conditions, generalizable -- instead of strain and/or environment-specific -- profiles are assembled that map the phenotypic and transcriptomic stress responses.

**Table 1.** Summary of the three *S. pneumoniae* strains in this study

| Strains | TIGR4 | Taiwan-19F | D39 |
| --- | --- | --- | --- |
| RefSeq | NC_003028 | NC_012469 | NC_008533 |
| Genome size (bp) | 2,160,842 | 2,112,148 | 2,046,115 |
| Number of genes | 2287 | 2119 | 2165 |
| Origin | Isolated from the blood of a<br>30-year-old male patient in<br>Kongsvinger, Norway | A patient with invasive<br>pneumococcal disease in<br>Taiwan | Clinical isolate, 1916 |
| Reference | Aaberge *et al.* (1995)<br>Tettelin *et al.* (2001) | Shi *et al.* (1995)<br>McGee *et al.* (2001) | Lanie *et al.* (2006) |
| Doubling time (min) |  |  |  |
| SDMM | 51 ± 5 | 74 ± 2 | 57 ± 4 |
| CDM | 80 ± 7 | 110 ± 10 | 74 ± 4 |
| MMCDM | 108 ± 9 | 118 ± 13 | 100 ± 8 |

### Genome-wide fitness and expression profiling reveal the phenotypic and transcriptional importance of cellular processes upon nutrient depletion

We applied two high-throughput, genome-wide methods to determine on two different levels how *S. pneumoniae* deals with nutrient depletion stress. The first level is measured by Tn-Seq, which quantifies a gene's effect on the growth rate (fitness) resulting from a genetic disruption by transposon insertions [18]. The second level is determined by RNA-Seq, which measures gene expression by quantifying transcript abundance. Tn-Seq fitness data thus provides insight into the phenotypic importance of each gene, while RNA-Seq provides insight into which genes respond to stress on a transcriptional level.

Six transposon insertion libraries for each strain were constructed and grown in each media to generate Tn-Seq profiles and a comprehensive genotype-phenotype map for each strain/environment pair (Figure 1a). As expected, and as we have shown before [1,2,14], the majority of insertions do not produce a significant fitness change, and more fitness defects (fitness < 1) were observed than increases in fitness (fitness > 1) (**Additional File 6**). To complement the Tn-Seq fitness data, genome-wide gene expression was measured for each strain/media pair by RNA-Seq. Four replicates from mid-exponential phase cells were prepared for each strain/media pair and sequenced at high depth (3.5-16 million reads/sample) [34]. Each sample contained reads mapping to 88-96% of all annotated genes in the corresponding genome, and transcript abundance between genes varied by nearly 10^5^ (Figure 1b, **Additional File 6**). Comparing the raw expression value (i.e. transcript abundance, not fold change) with fitness revealed that most genes with a fitness defect are highly expressed (Figure 1c), which is a pattern that holds across all strains and all three media. However, this pattern is of little predictive value, since the majority of highly expressed genes do not have a fitness defect or increase, and thus transcript abundance alone does not predict fitness.

**Figure 1:**
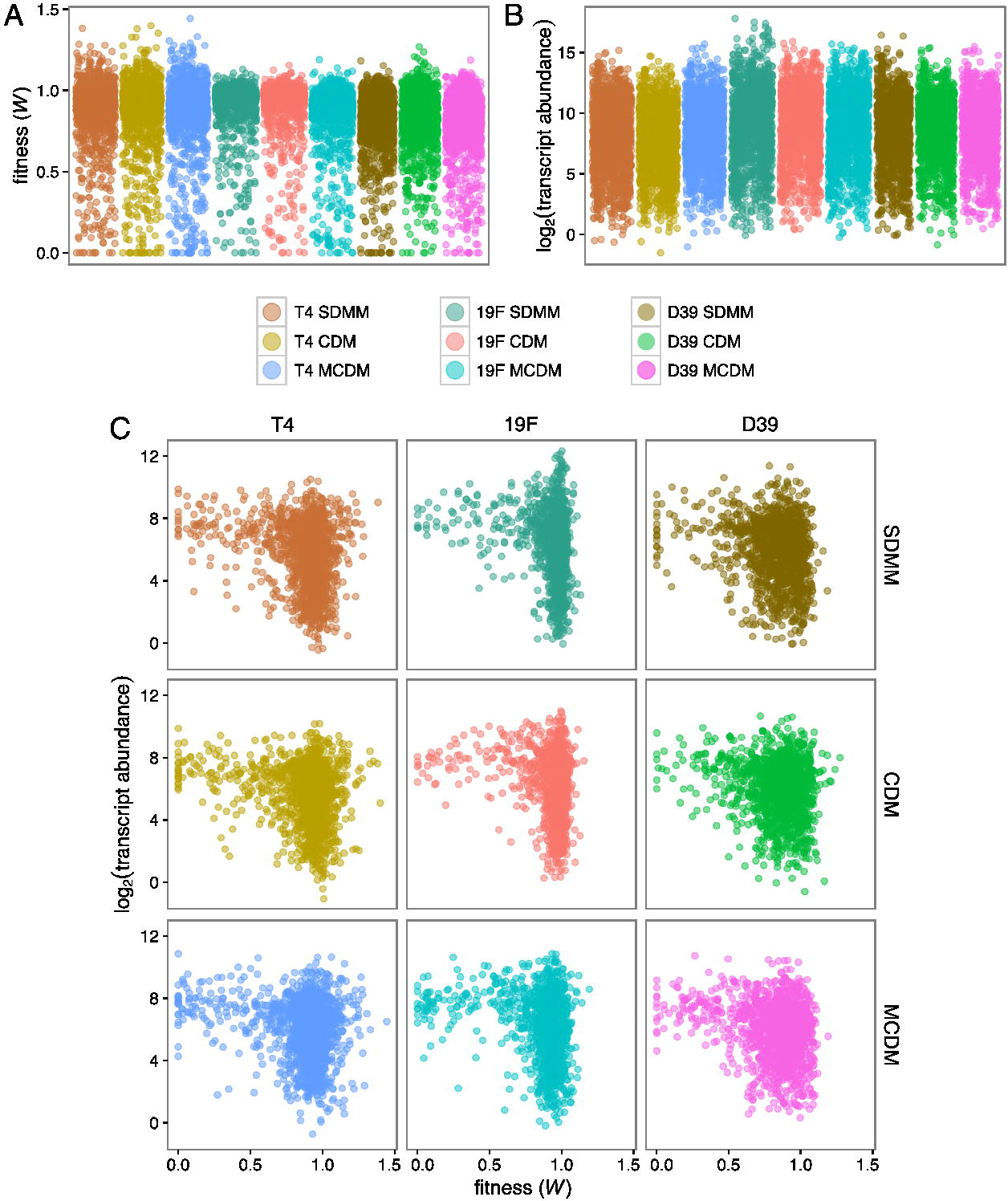
High-resolution profiles of phenotype and gene expression during stress incurred by nutrient depletion. **A**. Fitness values from Tn-Seq. **B**. Transcript abundance from RNA-Seq. Both high-throughput methods were performed on three *S. pneumoniae* strains (19F, T4, and D39) and in three media conditions (SDMM, CDM, and MCDM). **C**. By plotting Tn-Seq and RNA-Seq data on the same graph it becomes clear that genes with fitness defects (*W* < 0.85) are highly expressed.

### Validation experiments confirm that high-throughout profiles consist of high confidence data

The high-throughput genome-wide approaches Tn-Seq and RNA-Seq provide comprehensive profiles of cell physiology at two different levels. However, as with any high-throughput experiment the resulting datasets require careful validation to assess their accuracy. Tn-Seq data were validated by constructing 31 individual gene deletion mutants across the three strains. The mutants were used in monoculture growth assays and 1x1 competition assays in which the wild type strain is competed against the mutant to obtain fitness. In total 122 genotype-phenotype relationships were validated across the three strains and three media conditions, which to our knowledge is the largest validation set generated to date for different strains of a bacterial pathogen for which no ordered knockout arrays exist. This resulted in a strong correlation (R^2^ = 0.82), which is similar to correlations we achieved previously [1,2] and confirms high-confidence Tn-Seq fitness data (Figure 2a; **Additional File 7**). Importantly, these data also give detailed information on what type of stress is experienced by the bacterium. For example, Tn-Seq data shows that in rich media (SDMM) Δ*aroE (SP1376),* which catalyzes the conversion of 3-dehydroshikimate to shikimate and is associated with an upstream reaction in the *de novo* synthesis pathway of aromatic amino acids, has a growth defect (SDMM: *W* _*aroE*_ = 0.75), which we indeed validated (Figure 2b). However, a more severe defect in growth is measured when Δ*aroE* is grown in CDM and thus under stress by limited availability of amino acids (CDM: *W* _*aroE*_ < 0.26). Moreover, the gene becomes almost conditionally lethal in MCDM in the absence of aromatic amino acids (MCDM: *W* _*aroE*_ = 0.10). These data not only show that the Tn-Seq data consist of high confidence fitness values, but it also pinpoints environment-dependent weaknesses in the genomic network, highlighting how different genes become important in an environment-dependent manner. Moreover, it shows what type of stress is experienced in the environment; the conditional importance of *aroE* suggests an increasing lack of aromatic amino acids in CDM and MCDM.

**Figure 2:**
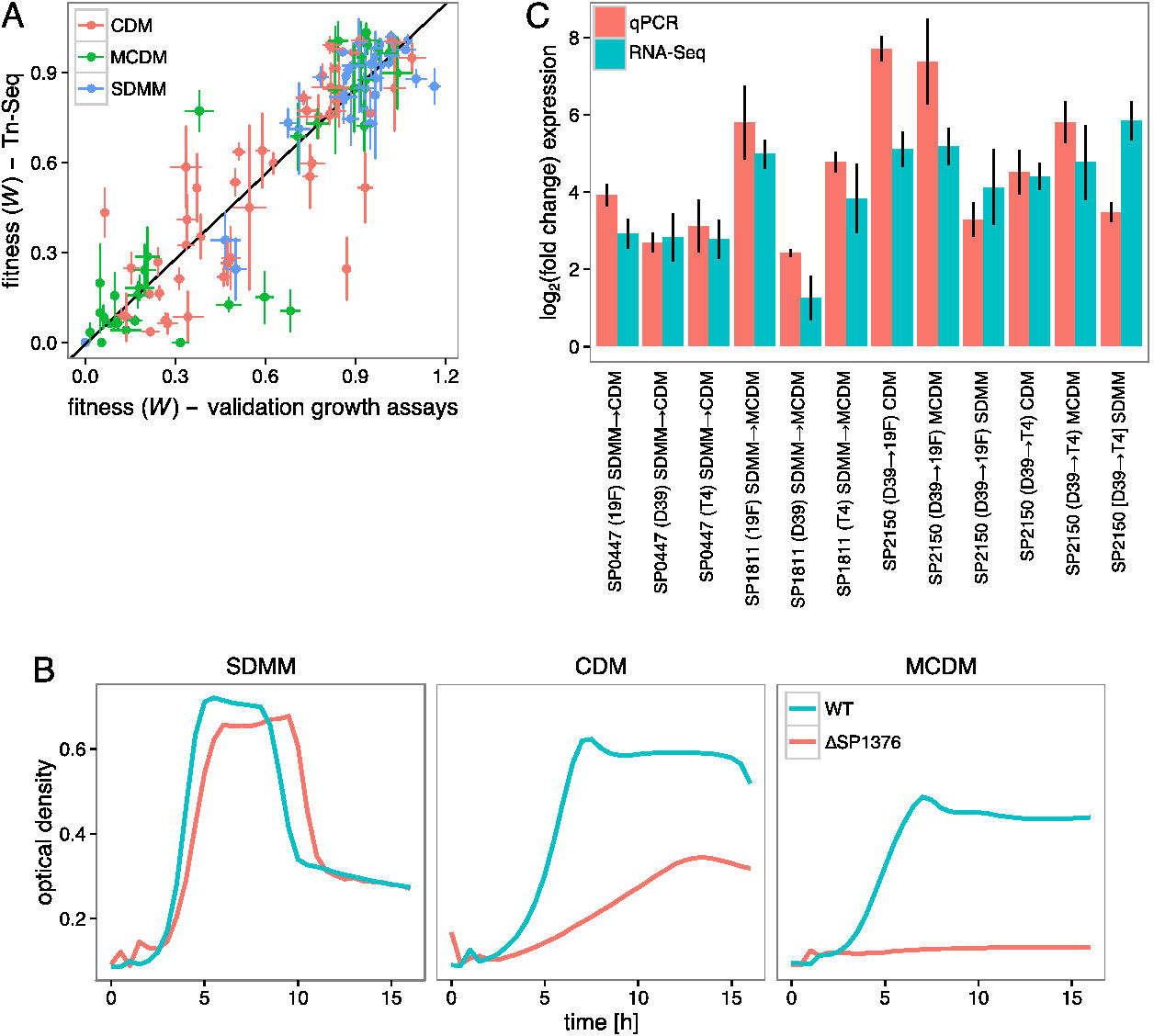
Tn-Seq and RNA-Seq data correlate strongly with validation experiments. **A**. A strong correlation between Tn-Seq fitness and fitness calculated from individual mutant growth curves or 1x1 competitions (*W* _*validation*_; n = 122; shown are mean ± SEM; linear fit yields R^2^ > 0.82) emphasizes that the strain-dependent sensitivity profiles are composed of high-confidence Tn-Seq data. **B.** Representative growth curves for comparisons between wild type (WT) and mutant ΔSP1376 in three different conditions, showing the dependence of the mutant on the presence of aromatic amino acids in the environment. **C**. Fold change in transcript abundance measured by qPCR and RNA-Seq. Vertical bars indicate SEM for both experiments. No qPCR and RNA-Seq pairs differed significantly (*p* < 0.0042, *t* -test with Bonferroni correction) indicating RNA-Seq data are composed of high-confidence data as well.

Additionally, the RNA-Seq data were validated by qPCR by measuring the expression of nine genes in the three media conditions (Figure 2c). The changes in expression measured by qPCR match the RNA-Seq differential expression data across all strains and media (**Additional File 8**), confirming that the generated genome-wide expression profiles represent real changes in transcription.

### Identifying genetic drivers of the nutrient stress response

The validated high-throughput data were used to identify genes responsible for the phenotypic (Tn-Seq) and transcriptomic (RNA-Seq) stress responses. By comparing fitness across environments we determined which genes changed their fitness (Δfitness, or Δ*W*) as *S. pneumoniae* transitions from rich media (SDMM) to defined (CDM) or minimal (MCDM) media (Figure 3a). Thus genes whose importance increases upon nutrient depletion will have a decreased fitness (*i.e.* a negative Δfitness) in the more restrictive media, while those whose importance does not change will have a Δfitness of 0. Those genes whose importance decreases will have an increased fitness (*i.e*. a positive Δfitness). Following a change from SDMM to either CDM or MCDM, each strain had an average of 12 genes increase in fitness and 29 genes decrease in fitness. For example, gene *SP1555* (dihydrodipicolinate reductase) is a key enzyme for lysine biosynthesis. A deletion of *SP1555* thus blocks *de novo* lysine synthesis and should hamper *S. pneumoniae*’s ability to overcome the depletion of extracellular lysine in CDM and MCDM. Indeed, Tn-Seq data shows that *SP1555* is not important in rich media (SDMM: *W_SP1555_* = 0.95), but shows decreasing fitness, and thus increasing importance, in the more stringent media (CDM: *W_SP1555_* = 0.78, Δ*W_SP155_* = - 0.17; MCDM: *W_SP155_* = 0.07, Δ*W_SP155_* = - 0.71).

**Figure 3:**
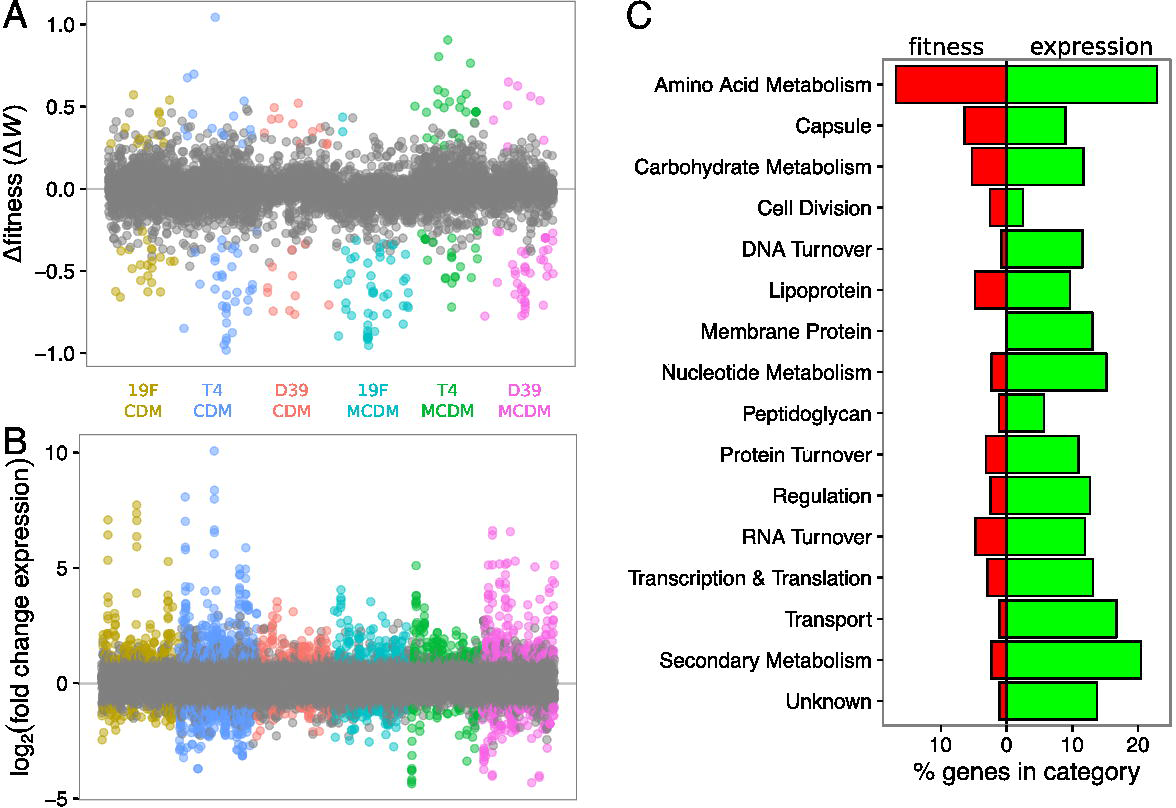
Changes in gene fitness and expression occur across all strains, media, and cellular subsystems. **A**. Δ*fitness* (Δ*W*) depicts how genes change their fitness as a strain transitions from rich media (SDMM) to defined (CDM) or minimal (MCDM) media. **B**. Shown are how genes change their expression as a strain transitions from rich media (SDMM) to defined (CDM) or minimal (MCDM) media. Expression is log_2_ fold change in transcript abundance from RNA-Seq. In both figures statistically significant changes are colored and both assays were performed on three *S. pneumoniae* strains (T4, 19F, and D39) and two media transitions (SDMM→CDM, SDMM→MCDM). **C**. Percentage of genes in each category with significant changes in fitness (red) and expression (green). For total number of genes in each category (by strain), see **Additional File 3**.

Besides genes that change in fitness, genes that change in expression were identified by comparing transcript abundances between SDMM and either CDM or MCDM (Figure 3b). On average, the media shift caused 101 genes to significantly increase expression and 125 genes to decrease expression. Overall, 5.4 times more genes showed significant expression changes compared to significant fitness changes. Importantly, in both the Tn-Seq and RNA-Seq datasets, the significant changes were distributed across a variety of cellular subsystems, indicating that the nutrient depletion environments trigger stress that is experienced network-wide (Figure 3c). Among metabolic subsystems, amino acid pathways are especially well represented, with 17% of genes showing a fitness change and 23% of genes differentially expressed (**Additional File 6**).

### Genome-wide data visualization with a whole cell model reveals that expression profiles are poor predictors of phenotypic importance

*S. pneumoniae* designates a large fraction of its genome to metabolism, and a number of metabolic enzymes have been linked to the bacterium's phenotypic stress response to nutrient and antibiotic perturbations [1,2]. Given the large number of metabolic genes with fitness or expression changes during nutrient depletion (Figure 3c), we focused on metabolic pathways to identify patterns in how *S. pneumoniae* handles and overcomes stress. To analyze metabolism in a systematic and comprehensive way, a genome-scale metabolic model of *S. pneumoniae* was assembled. A draft metabolic model was derived from the KBase system [35] (http://kbase.us) by collecting metabolic reactions associated with the annotated genomes for T4, 19F, and D39. To account for non-enzymatic reactions and reactions with misannotations, a gap-filling algorithm [36] added reactions to ensure growth of the model on SDMM, CDM, and MCDM. Pathways in the model were manually curated by comparing reactions and gene associations to KEGG [37] and BioCyc [38], with a particular emphasis on amino acid and nucleotide metabolism (**Additional Files 9 and 10**). The final model, called iSP16, details the interconversion of 866 metabolites by 928 reactions, catalyzed by 463 genes (43.9% of all ORFs in T4). To our knowledge, iSP16 is the first curated, genome-scale metabolic model in the genus *Streptococcus*.

Using iSP16, we searched for patterns in the location of fitness and expression changes during nutrient stress. Initially, we expected the Tn-Seq and RNA-Seq data to align, as increasing expression of a metabolic enzyme can increase flux through an important pathway [39]. However, overlaying the datasets onto the network shows that, when transitioning from SDMM to CDM, metabolic pathways either change on a transcriptional level or their fitness (phenotypic importance) changes, but they almost never change in the same location (Figure 4a). Moreover, upon transitioning to MCDM, the clusters of fitness or expression changes expand but they almost never merge (Figure 4b). This means that, contrary to our expectations (and popular belief), overlap between fitness and expression changes are incredibly rare (Figure 4).

**Figure 4:**
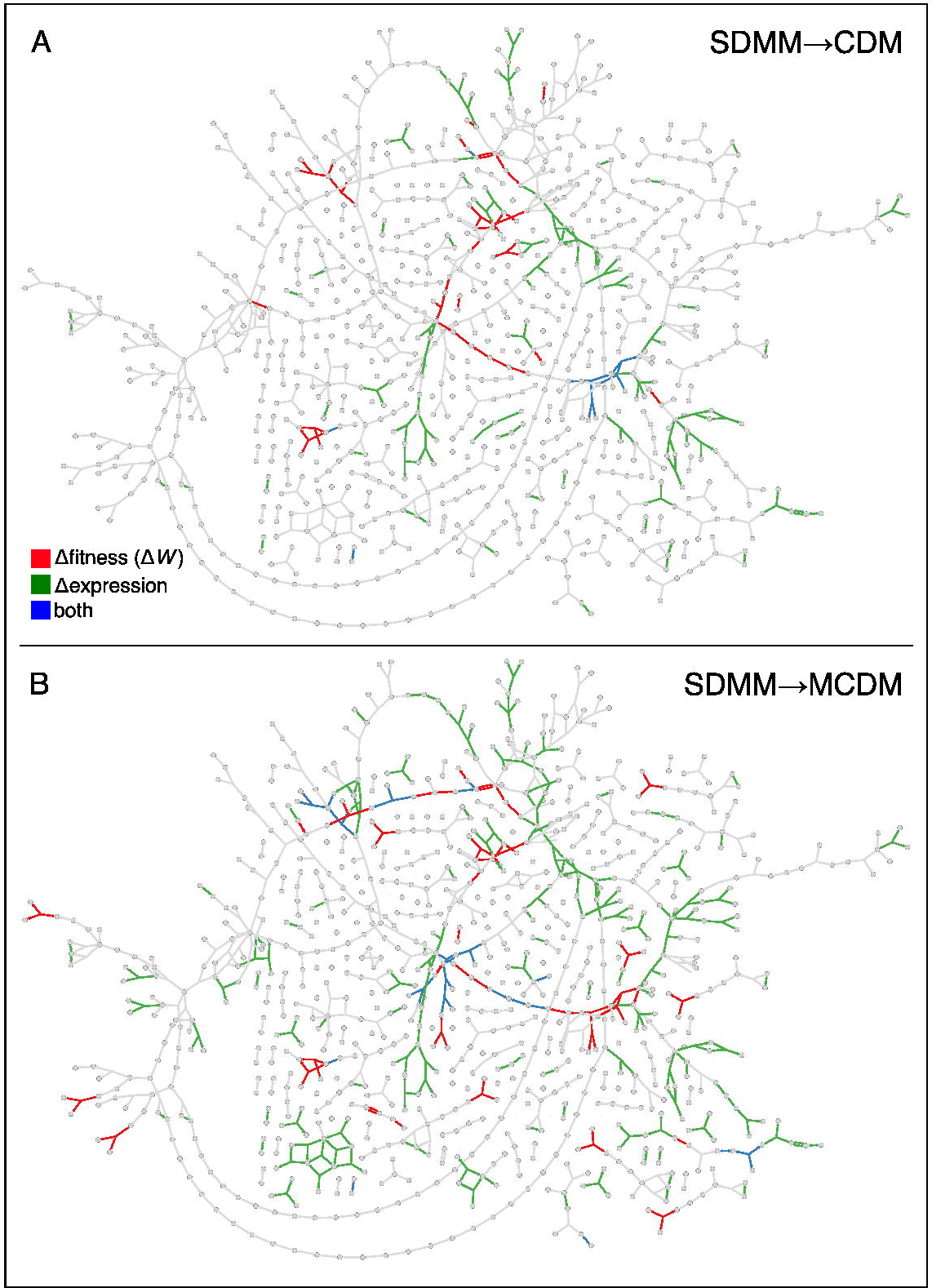
Genes with significant changes in expression (green), fitness (red), or both (blue) are distributed throughout the iSP16 metabolic model. Lines indicate reactions connecting metabolites (circles). Minor and currency metabolites are not shown (see **Additional File 9**). Reactions are colored based on gene associations in the iSP16 model.

A clear example of the disconnect between fitness and expression changes is the shikimate pathway, the biosynthetic route for the aromatic amino acids (Figure 5a). Beginning with the glycolytic and pentose phosphate pathway intermediates phosphoenol-pyruvate (PEP) and erythrose-4-phosphate (E4P), the shikimate pathway uses seven enzymatic reactions before branching in sub-pathways specific for tryptophan (Trp), phenylalanine, and tyrosine. The branchpoint occurs immediately after gene *SP1374*, with gene *SP1816* catalyzing the first reaction of the Trp-specific branch (Figure 5a). Compared to SDMM, CDM contains reduced tryptophan, and MCDM contains no tryptophan (Table 2, **Additional File 4**). We expected that the removal of Trp would cause increased expression of the shikimate pathway to compensate for the absence of Trp and decreased fitness when any of the pathway's genes are interrupted. Although this pathway shows both fitness and expression changes in our data, the fitness changes are restricted to the genes above the branch into tryptophan synthesis (*SP1700* - *SP1374*), while the expression changes are below the branch point (*SP1817* - *SP1812*) (Figure 5a). The sole biosynthetic route to tryptophan thus contains disjoint sets of genes with fitness and expression changes, indicating that a single pathway can be split between the phenotypic and transcriptional stress responses.

**Table 2.**
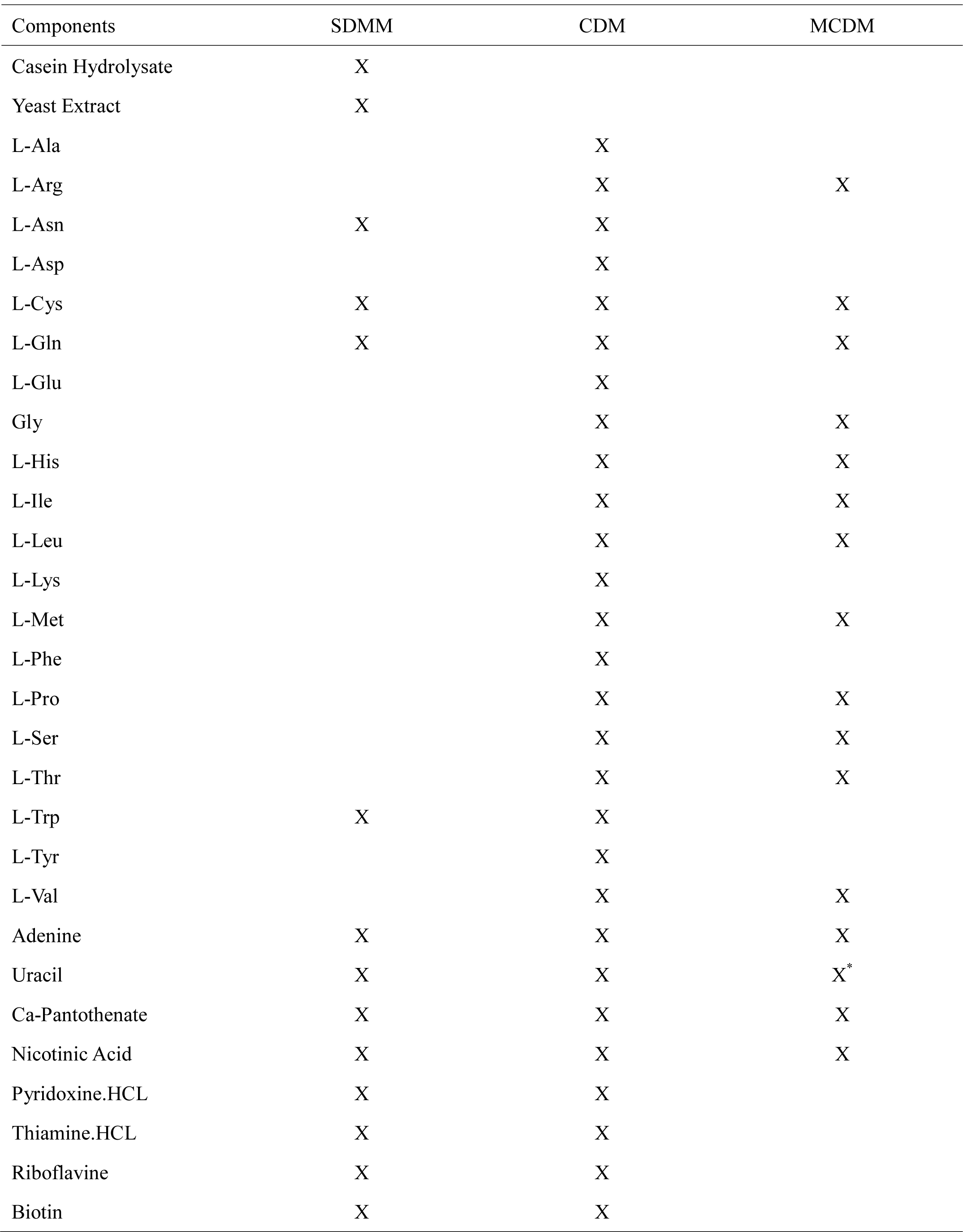
Comparison of nutrient availability in SDMM, CDM and MCDM.

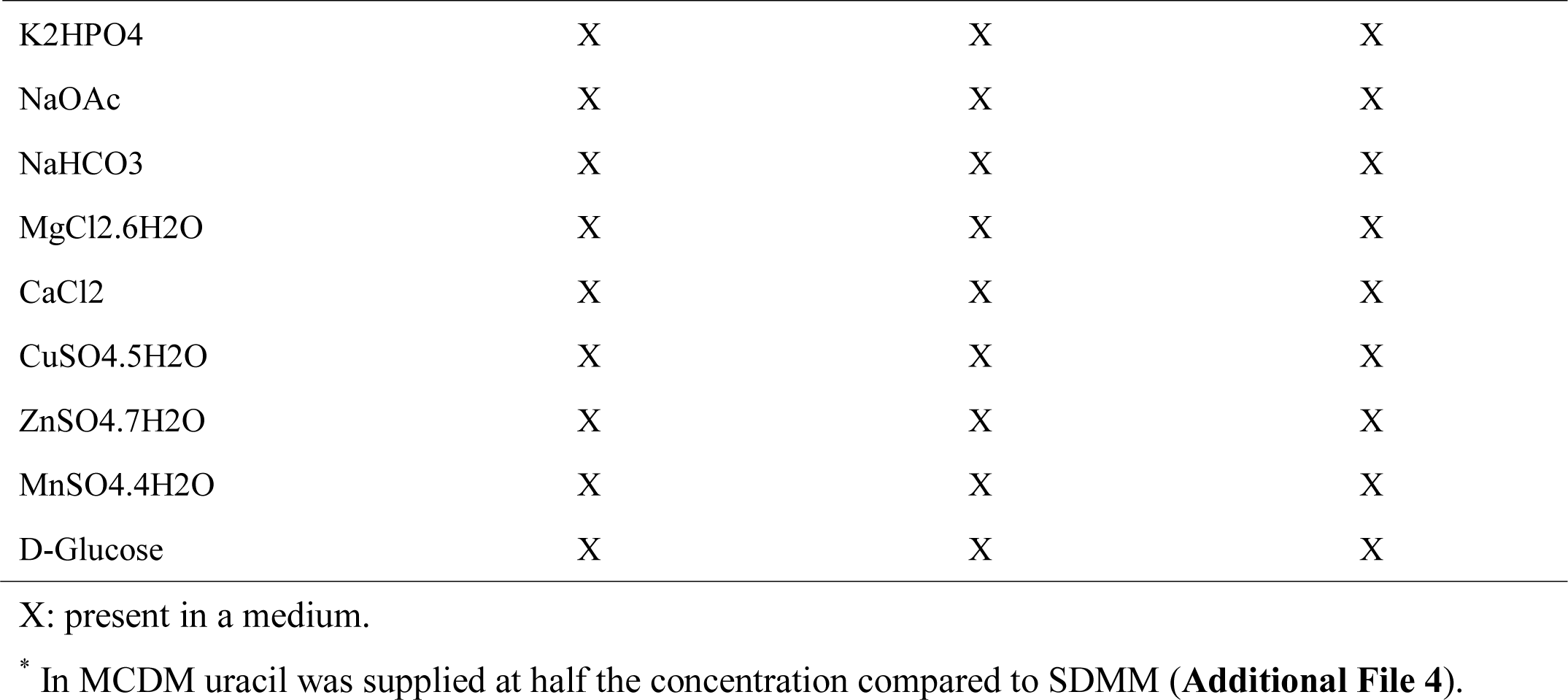

**Figure 5:**
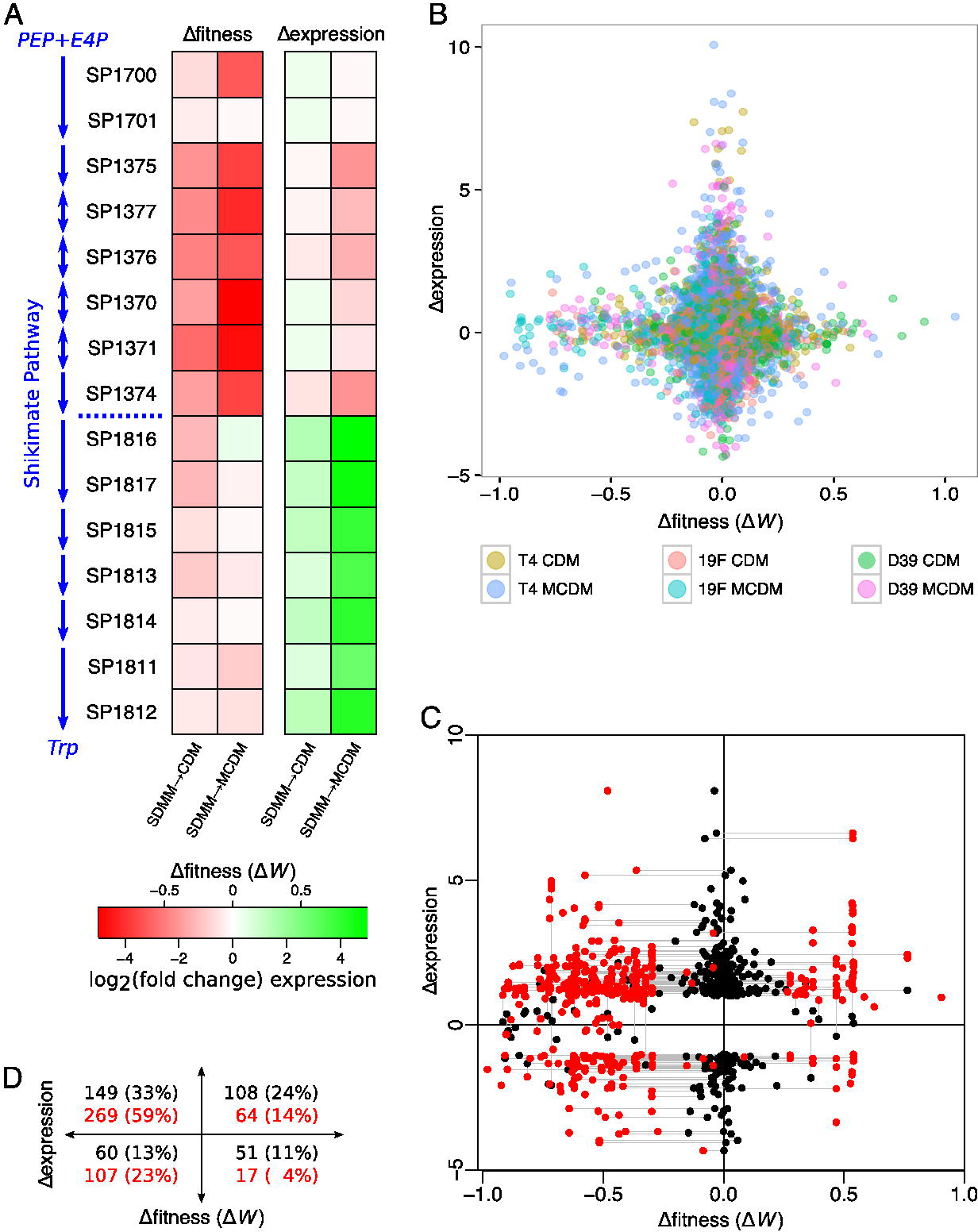
Fitness and expression changes do not occur on the same gene but appear to be related. **A**. Changes in gene expression and fitness are separated in the tryptophan biosynthesis branch of the shikimate pathway. Single and double-headed arrows indicate reversibility or non-reversibility of individual chemical reactions, respectively, while arrows spanning two genes indicate enzymatic subunits catalyzing the same reaction. The dashed blue line (between SP1374 and SP1816) indicates the branch point into tryptophan biosynthesis. PEP=phosphoenol-pyruvate, E4P=*D* -erythrose-4-phosphate, Trp=*L* -tryptophan. **B**. Changes in gene expression (RNA-Seq: Δ*expression*) vs. changes in fitness (Tn-Seq: Δ*fitness* (Δ*W*)) are not correlated. For each data point (gene) Δ*fitness* (Δ*W*) and Δ*expression* are plotted depicting how a gene’s fitness and expression change as a strain transitions from rich media (SDMM) to defined (CDM) or minimal (MCDM) media. **C**. Genes with significant Δ*expression* or Δ*fitness* (Δ*W*) (black) migrate off the horizontal and vertical axes when paired with the nearest neighbor that changes in fitness or expression (red) (NB. If multiple genes appear at the same distance from the gene with a fitness change, the expression changes are averaged.). **D**. The number of genes in each quadrant of the Δ*expression/* Δ*fitness* plot shifts before (black) and after (red) nearest-neighbor pairing, showing that fitness and expression changes are somehow linked.

Strikingly, the lack of correlation between fitness and expression changes is not limited to metabolic genes. When plotting genome-wide expression changes against fitness changes (Figure 5b), no clear relationship is observed between fitness and expression changes; almost all genes appear on either the horizontal or vertical axes of Figure 5b, indicating either a fitness change with no expression change, or vice versa. Therefore, even though fitness and expression changes occur across the genome, their disjointedness would go unnoticed if only Tn-Seq or RNA-Seq experiments were performed, suggesting that upon exposure to stress, expression profiles are not good predictors of what is actually phenotypically important in a bacterium.

### Cellular networks link changes in gene expression and fitness

In the metabolic network, fitness and expression changes rarely overlap, but they often seem to be located near one another (Figure 4). Thus even though expression changes are not indicative of genes that are phenotypically important, expression changes suggest that phenotypically important genes are often very close neighbors. To test this hypothesis a nearest neighbor analysis was performed in which each gene with a fitness change is paired with the closest gene in the network with an expression change (Figure 5c). The same procedure was repeated, starting with the genes with expression changes and searching for nearby fitness changes. Importantly, the nearest neighbor transformation restores the expected relationship between increased expression and decreased fitness. Prior to considering the nearest neighbors, 33% of the genes in Figure 5a appeared in the upper left quadrant (decreased fitness and increased expression) (Figure 5d). After pairing with nearby genes, 59% of the genes moved into this quadrant, the largest change for any quadrant of the graph.

This raises at least two questions: 1.) How close, on average, are paired neighbors; and 2.) Is there a neighbor-to-neighbor relationship between the size of the fitness change and the change in expression? For instance, are genes with the largest fitness changes neighbors with genes with the largest expression changes? To answer these questions distances between genes in the metabolic network were quantified by counting the number of reactions between enzymes (Figure 6a). For instance, genes associated with the same reaction have a distance of zero, while genes associated with reactions that share a common metabolite have a distance of one. To quantify the magnitude of both the fitness and expression changes, we multiply the two changes. This product corresponds to the area of a rectangle drawn between a point in Figure 5c and the origin (Figure 6b). This “area off axes” is maximized when both the fitness and expression changes are large, and it is near zero when either the fitness or expression change is small. Finally, by plotting the fitness-expression product against the distance between the corresponding genes (Figure 6c), two conclusions can be drawn. First, the majority of fitness and expression changes pair at short distances, with 80% of all pairs within a distance of two, and 93% within a distance of three. Second, the size of the fitness and expression changes in each pair decreases with distance (*p* < 0.005, ANOVA). This analysis (Figure 5) thus shows quantitatively what is visually suggested in Figure 4; fitness and expression changes occur in distinct, but co-localized genes. Moreover, the largest changes in fitness are close to the largest expression changes, which means that fitness and expression changes are not only co-located, but have a magnitude that matches their neighbors'.

**Figure 6:**
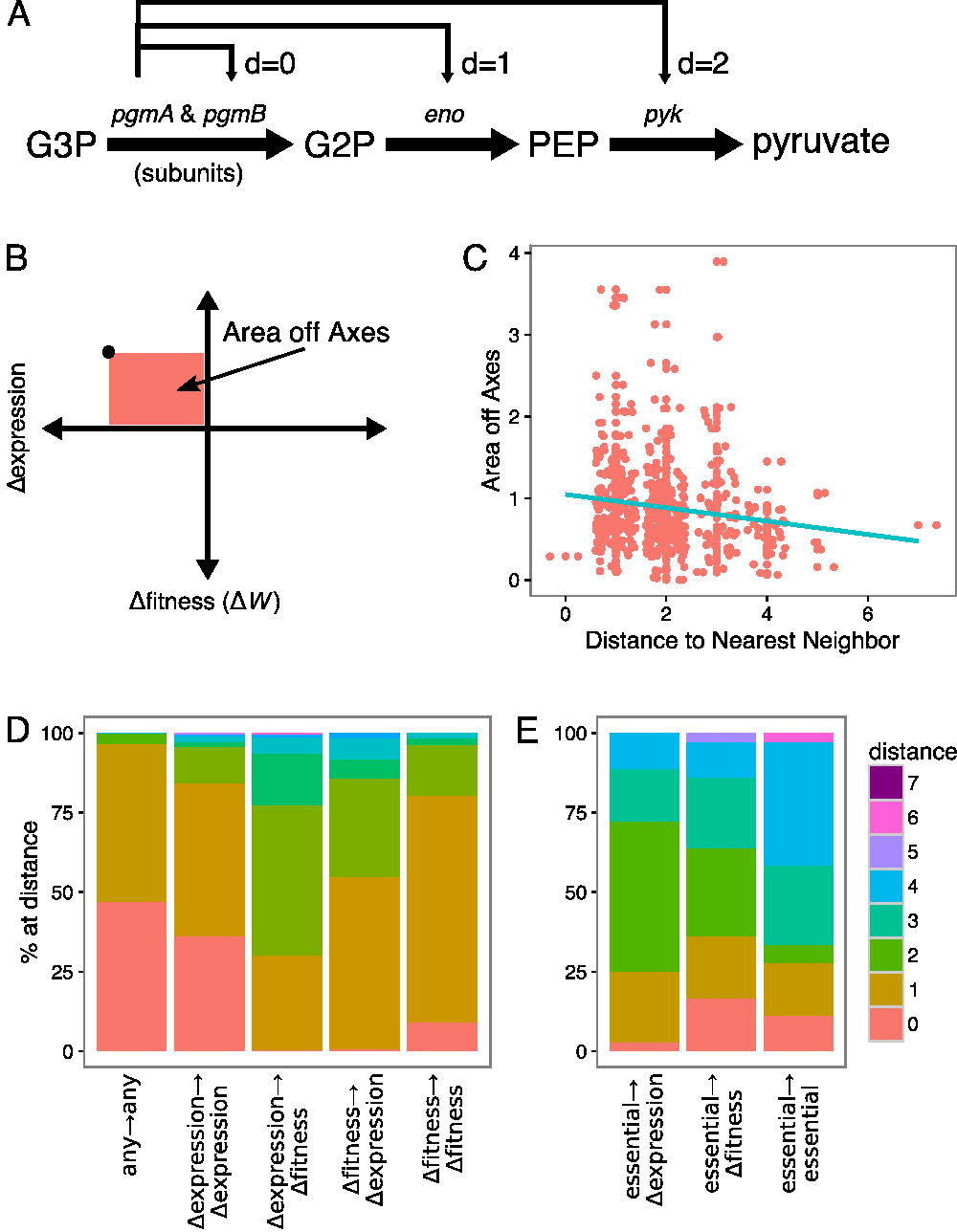
Metabolic models can quantify distances between reactions and genes. **A**. Shown is how distances are calculated. For instance, the distance between the subunits *pgmA* and *pgmB* in glycolysis is zero, while either enzyme has a distance one to *eno* and a distance two to *pyk*. **B**. The area off axes is the product of Δ*expression* and Δ*fitness* (Δ*W*) and is maximized when both the fitness and expression changes are large. The area off axes is near zero when either the fitness or expression change is small. **C**. Extent of off-axis migration after nearest-neighbor averaging decays with distance between the gene and its nearest neighbor, indicating that changes are closely located within the network and that the largest changes in fitness are close to the largest expression changes. **D**. Distribution of network distances varies between genes with fitness and expression changes. Distances between all genes (*any*), genes with significant changes in expression (Δ*expression*) or fitness (Δ*fitness*). Unconnected genes (distance = ∞) are not shown. The short distances between two genes with expression changes or two genes with fitness changes indicates small subnetworks of either fitness or expression changes. **E**. Essential genes and genes with fitness or expression changes are not evenly distributed. On average, essential genes (*essential*) are closest to genes with fitness changes (Δ*fitness*) and farthest from genes with expression changes (Δ*expression*).

Further visual inspection of the metabolic network maps (Figure 4) suggests that genes with fitness changes and genes with expression changes are not distributed evenly throughout the network. Instead, it appears that small clusters of fitness and expression changes are present, suggesting small sub-networks that either change transcriptionally or that are conditionally important. To quantify the existence of these clusters, we compared the distribution of distances among all genes, genes with fitness changes, and genes with expression changes (Figure 6d). Genes with an expression change (Δexpression→Δexpression) are indeed clustered at shorter distances to other differentially expressed genes, and the same is true for genes that change in fitness (Δfitness→Δfitness) (*p* < 0.005, *t* -test of Poisson fit to distributions of distances) (Figure 6d). In contrast, essential genes, derived from these and previous Tn-Seq data [2,14] (**Additional File 11**) are not closely clustered (Figure 6e, essential→essential). Instead, essential genes are closer to genes with fitness changes than they are to either genes with expression changes or other essential genes (Figure 6e, essential→Δessential and essential→Δfitness, *p* < 0.05). This distribution shows that: 1.) Phenotypic stress response genes lie closer to essential genes than transcriptional stress response clusters; and 2.) Both transcriptional control and important functions are intertwined, but separated into small clusters that correspond to either the phenotypic or transcriptional stress responses. Therefore, it appears that genes that become phenotypically important in a new environment are shielded from highly fluctuating changes in expression, while nearby genes that are less important on a phenotypic level may fluctuate to a much larger extent, suggesting these different sets of genes are part of differently organized regulatory modules.

### Meta-analysis reveals partitioning of phenotypic and transcriptomic stress responses across multiple conditions

If these distinct sets of genes are indeed organized in different regulatory modules, this would suggest that essential genes and the phenotypically important genes we identified would be universally shielded from expression changes across any condition. To test this hypothesis, we performed a meta-analysis using all publicly available *S. pneumoniae* datasets in the Gene Expression Omnibus (GEO) database. Using microarray data from 234 experiments (**Additional File 9**), we calculated the “expression plasticity” across a wide range of genetic and environmental perturbations (Figure 7a). The plasticity is the relative variance in expression of the gene (and its homologs) across all datasets in the GEO database. This means that genes with low plasticity rarely vary in their expression levels, while high plasticity genes vary widely in their expression across conditions. The GEO meta-analysis shows that both essential genes (which are phenotypically important in all environments) as well as genes with fitness changes in our dataset have low expression plasticity across the GEO datasets (Figure 7b). Furthermore, plotting the GEO expression plasticity against the size of the fitness change in our experiment reveals that genes with the largest fitness changes have the lowest plasticity (Figure 7c). And thus, not only do the phenotypically important genes have no corresponding expression change in our experiments, they also hold their expression relatively constant across all the experiments in GEO. This meta-analysis suggests that the partitioning of the phenotypic and transcriptional stress responses extends beyond the nutrient depletion stresses analyzed in this study and further suggests that the phenotypic and the transcriptional response networks are composed of highly dissimilar regulatory modules.

**Figure 7:**
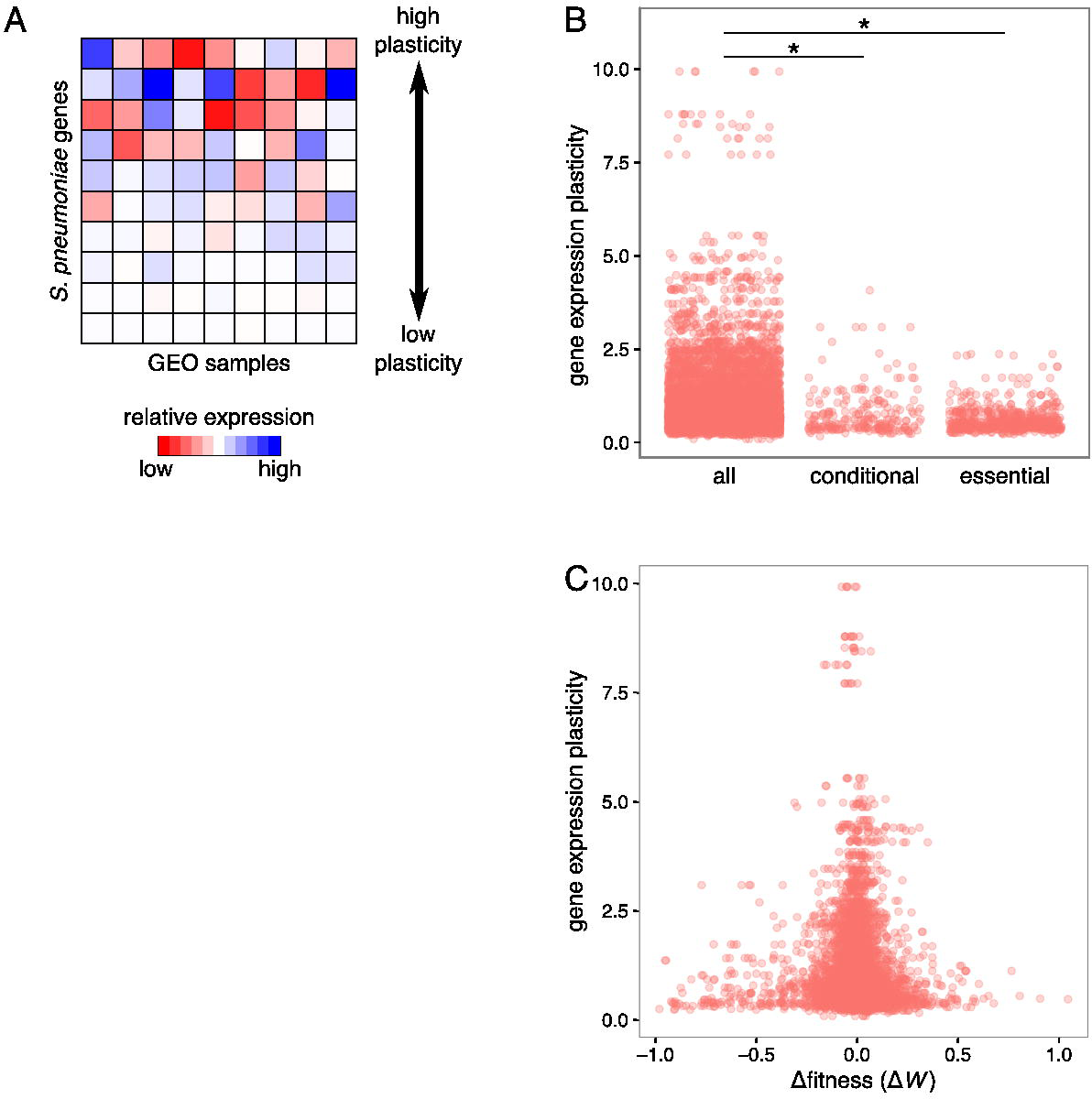
Meta-analysis shows that essential and phenotypically important genes are shielded from large changes in expression. **A**. Quantifying gene expression plasticity using meta-analysis of GEO expression studies. Plasticity is the normalized variance in gene expression across all *S. pneumoniae* data in GEO (see **Additional File 9**). **B**. Gene expression plasticity is significantly lower for essential genes (*p* < 10^−14^) and conditionally essential genes (genes with a significant fitness change) (*p* < 10^−34^, both comparisons *t* -test on means of fitted Gamma distributions). Both the essential and conditionally essential genes appear to be shielded from transcriptional changes not only in our RNA-Seq data, but across all the expression datasets in GEO. **C**. Gene expression plasticity decreases with increasing magnitude of the fitness change (relative to SDMM). Thus the amount of shielding is proportional to a gene's phenotypic importance, and genes with the largest fitness changes show the smallest variation in expression across the experiments in GEO.

## CONCLUSIONS

A popular assumption is that differentially expressed genes are good proxies for genes that are important for maintaining the survival of a microbe under stressful conditions [8-11]. Our validated genome-wide profiles for three strains of *S. pneumoniae* show in detail that genes and pathways that become important upon nutrient depletion are generally not differentially expressed. It is possible that the products of the genes that change their fitness are characterized by post-transcriptional or post-translational, rather than transcriptional, changes [40,41]. For instance, enzymes involved in protein and carbohydrate biosynthesis are heavily modified by lysine succinylation in multi-drug resistant *Mycobacterium tuberculosis* strains [42], while S-glutathionylation serves as a major regulation mechanism when *Cyanobacteria* are under oxidative stress conditions [43]. However, in general these mechanisms only affect a relatively small number of genes, and overall there seems to be a strong correlation between transcription and protein levels, especially during mid-exponential growth [44]. We thus do not believe that our findings are coincidental and related to our organism or experimental setup, also because several other studies on bacterial and fungal species have implied similar patterns in certain metabolic pathways and cellular functions, even though not all of them were validated or conducted on a genome-wide level [40,45,46]. It seems that a poor correlation between transcriptional change and functional importance is, at least for bacteria, universal. This means that transcriptomic profiling studies should be assessed carefully when they are used as a predictor of genes that matter phenotypically, e.g. in genetic engineering or drug target identification studies. Drug target identification across different domains of life has heavily relied on transcriptomic data as a surrogate for functional importance [47-53]. For instance, inhibitors of streptokinase gene expression have been proposed as novel antimicrobials for group A streptococcus [52], and gene expression datasets were used as a source for co-target identification using a random-walk model based on *M. tuberculosis* [51]. Our results provide an important argument to search for phenotypically important genes instead of differentially expressed genes in future antimicrobial discovery -- blocking differentially expressed genes may fail to cause a growth defect while phenotypically important genes directly affect an organism’s fitness.

In our attempts to unify fitness and expression changes, we showed that leveraging genome-scale metabolic modeling and topological analyses links both sets of genes in a cellular metabolic network. Although changes in transcription status and changes in functional importance occur on separate sets of genes, our result show that these sets of genes are either in the same pathways or closely related pathways that share intermediates. This pattern is consistent with the ideas of metabolic control analysis (MCA), where pathway flux can be controlled by changes in either enzyme abundance or substrate concentrations [39,54]. A central result of MCA is that the relative importance of enzyme or substrate changes can vary along a pathway. In our example of the shikimate pathway (Figure 5a), the upper half of the pathway (SP1371-SP1377) may be controlled by substrate abundance, while flux through the amino acid-specific branches (e.g. SP1811-SP1818) may be controlled by enzyme abundance. It is important to note that all of the reversible reactions in the shikimate pathway occur before the branch point (Figure 5a), and the (reversible) enzymes above the branch point are not differentially expressed. Reversibility allows pathway intermediates to control flux through feedback mechanisms, possibly lessening the importance of transcriptional control in the upper branch. By deferring most of the transcriptional control until the lower branches of the shikimate pathway, *S. pneumoniae* may be able to control the production of Trp separately from the other aromatic amino acids while still maintaining adequate flux through the upper branch via substrate-level feedback.

A common explanation for the lack of an expected fitness defect is redundancy in the surrounding network. Since all of the fitness defects in the shikimate pathway occur before it branches into pathways for individual amino acids, it is possible that redundancies exist that can overcome the loss of a biosynthetic route for a single amino acid, but not for all three. Although there is no known alternative route for aromatic amino acid synthesis outside of the shikimate pathway, the Trp-specific genes could be redundant and thus some genes in the pathway could alleviate the absence of other genes and provide functional redundancy. Identifying these “hidden” redundancies is a powerful benefit of overlaying genome-scale phenotypic data onto mathematical models.

In addition to allowing more parsimonious gene regulatory networks, separating transcriptional control from phenotypic importance may allow bacteria more flexibility to respond to new environments without incurring a fitness cost. Genes that fluctuate transcriptionally are not important for sustaining growth and hence have more flexibility in their expression. Phenotypically important genes seem to be shielded from large, fluctuating expression changes and are possibly controlled by feedback loops, which is a mechanism adept at retaining expression levels within tight boundaries [55]. Taken together, these features allow a bacterium to maintain a tightly controlled, robust core of essential genes while simultaneously preserving metabolic flexibility.

In this report, we present a transferable, systems-level approach to reconcile transcription and fitness changes within a network, which serves as an important attempt to achieve a systems-level understanding of how a bacterium deals with environmental perturbations. Although metabolism represents the majority of genes that change in either expression or fitness during nutrient depletion (Figure 3c), we are striving to achieve a truly holistic view by integrating additional parts of the genomic network, including energy generation, cell division, transport, and formation and turnover of the cell wall and membrane. Lastly, applying network topological analyses to contextualize high-throughput experiments has the potential to provide value in genetic engineering, predicting drug target candidates, and re-evaluating current drug targets with the goal to achieve a higher success rate in developing novel strategies to eradicate microbial pathogens.

## METHODS

### Bacterial strains, growth and media

Experiments were performed with *S. pneumoniae* strains TIGR4 (T4; NCBI Reference Sequence: NC_003028.3) and Taiwan-19F (19F; NC_012469.1), and D39 (NC_008533). All gene numbers are according to the TIGR4 genome, except the unique genes, which are preceded by SP, SPT, and SPD for TIGR4, Taiwan-19F, and D39, respectively. A "correspondence table" that matches homologous genes in the three strains can be found in **Additional File 2**. Single gene knock-out strains were constructed by replacing the coding regions with a chloramphenicol or spectinomycin resistance cassette as described previously [2,14,56]. Except for Tn-Seq experiments, RNA-Seq experiments, and specific growth conditions, *S. pneumoniae* was cultivated statically in Todd Hewitt broth with 5% yeast extract and 5 µL/mL of Oxyrase (Oxyrase, Inc), or on sheep’s blood agar plates at 37°C in a 5% CO 2 atmosphere. When appropriate, liquid culture and blood agar plates contained 4 µg/mL of chloramphenicol (Cm) or 200 µg/mL of spectinomycin (Spec) for selecting strains or mutant libraries that contain drug markers. Tn-Seq and RNA-Seq experiments were performed in three growth media that contain gradually decreasing nutrient levels, namely, semi-defined minimal medium (SDMM) [2], chemically defined medium (CDM; **Additional File 4**) and minimal chemically defined medium (MCDM; **Additional File 4**).

### Tn-Seq library construction and selection experiments

Transposon insertion mutant libraries were constructed as previously described with the transposon Magellan6, which lacks transcriptional terminators therefore allows for read-through transcription and diminishing polar effects [2,14,57]. Additionally, the mini-transposon contains stop codons in all three frames in either orientation when inserted into a coding sequence. Six independent transposon libraries were constructed in T4, 19F, and D39. Tn-Seq experiments were performed with each transposon library for each of the three strains under the three media conditions (pH 7.3 with 20mM of supplemental glucose in SDMM and MCDM and 28 mM glucose in CDM). Mutant libraries were grown to mid- to late-log phase and harvested for genomic DNA isolation.

### Tn-Seq sample preparation, sequencing and fitness calculation

Sample preparation, Illumina sequencing, and fitness calculations were performed as previously described [2,14,18,58]. For each insertion, fitness (*W* _*i*_) representing the growth rate is calculated by determining the change in frequency for the mutant in the population [58]. Fitness for single genes is calculated by averaging *W* _*i*_ over all the insertions in the same gene. To determine whether fitness effects significantly differ between conditions, three requirements must be fulfilled: 1.) W_*i*_ is calculated from at least three data points (insertions), 2.) the difference in fitness between conditions has to be larger than 15% (|*W* _*i*_ - *W* _*j*_| > 0.15), and 3.) the difference in fitness has to be significantly different in a one sample *t* -test with Bonferroni correction for multiple testing.

### Competition and single strain growth assays

1x1 competition experiments were performed by mixing a single gene knock-out strain with the corresponding wild type strain in a 1:1 ratio and growing for approximately eight generations to late exponential phase in a particular growth medium [2]. A sample was taken at the beginning and the end of a competition experiment for CFU counts and plated on blood agar plates (for a total CFU count) and on blood agar plates with selective antibiotics (for a CFU count of the knock-out strain). Fitness was then calculated using the same approach as Tn-Seq by determining the ratios of the competing strains at the start and end of the competition and determining the expansion of the population using CFU counts. Single-strain growth assays were performed in 96-well plates by taking OD_600_ measurements on a Tecan Infinite 200 PRO plate reader. Both competition and single-strain growth assays were performed no fewer than three times.

### RNA-Seq sample collection

The three strains were grown under each of the three growth media conditions (SDMM, CDM, and MCDM) in four biological replicates. 5 mL of early mid-log phase liquid culture was harvested by centrifugation at 4°C, snap frozen, and stored at -80°C for RNA isolation. Total RNA was isolated using the RNeasy Mini kit (Qiagen).

### RNA-Seq sample preparation, sequencing and expression level calculations

RNA-Seq cDNA libraries were generated following the RNAtag-Seq protocol as previously described [59]. Briefly, 400ng of RNA was fragmented, depleted of genomic DNA using TURBO DNA-free kit (Ambion), 5’-dephosphorylated, and subsequently ligated to barcoded RNA adapters at the 3’-terminus. Barcoded RNA samples were pooled and purified with RiboZero (Illumina). The ribosomal RNA-depleted samples were converted to Illumina cDNA sequencing libraries in three key steps: 1.) First strand cDNA synthesis with AffinityScript Multiple Temperature cDNA Synthesis kit (Agilent) and RNA degradation, 2.) ligation to a 3’-linker, and 3.) PCR amplification using primers that target the 3’-linker and a constant region of the RNA barcodes and contain the Illumina flow cell sequences. The cDNA libraries were sequenced on an Illumina NextSeq500 platform (single read, 50 base pair).

Raw reads were demultiplexed, trimmed to 40 base pairs, and quality filtered using custom R scripts and the ShortRead package. Reads were mapped to the corresponding *S. pneumoniae* genome using Bowtie [60] with settings "-n 2 -l 60 -m 1 -B 1". Reads were aggregated to genes using the GenomicRanges R package and differential expression was calculated using DESeq2 [61].

### qPCR expression analysis

The three wildtype strains were grown in SDMM, CDM, and MCDM to early mid-log phase. Sample collection and total RNA isolation were performed following the same procedure as RNA-Seq. 4 ug of RNA from each sample was treated with the TURBO DNA-free kit, after which 400 ng of cleaned-up RNA was subjected to first strand cDNA synthesis using iScript reverse transcription Supermix (BioRad). Quantitative PCR was performed using a BioRad MyiQ; each sample was measured in two biological replicates and three technical replicates. No-reverse transcriptase and no-template controls were included for each sample. Expression levels from all samples were normalized against the 50S ribosomal gene SP2204.

### Statistical analysis

Statistical analyses were performed in R (http://www.r-project.org). Gene distributions were fit to either Poisson (gene distance distributions) or Gamma (GEO plasticity) distributions using the fitdistr function in the MASS toolbox. Expected values of the distributions were compared by a *t* -test.

## DECLARATIONS

**Ethics approval and consent to participate**. Not applicable.

**Consent for publication**. Not applicable.

**Availability of data and materials**. The datasets generated during the current study are available as Additional Files and in the Sequence Read Archive (SRP082544).

**Competing interests**. The authors declare that they have no competing interests.

**Funding**. This work was supported by the NIH R01 AI110724 and U01 AI124302.

**Authors' contributions**. PAJ led the computational experiments. ZZ led the wet-lab experiments. PAJ and ZZ performed computational and wet-lab experiments. All authors analyzed data, interpreted results, and wrote and approved the final manuscript.

**Acknowledgements**. DNA sequencing was performed at the Boston College Sequencing Core.

## ADDITIONAL FILES

**Additional File 1:** *S. pneumoniae* phylogenetic tree.

**Additional File 2:** Gene correspondence table.

**Additional File 3:** Breakdown of *S. pneuomniae* genes by strain and category.

**Additional File 4:** Media composition for SDMM, CDM, and MCDM.

**Additional File 5:** Growth curves for T4, 19F, and D39 in SDMM, CDM, and MCDM.

**Additional File 6:** Tn-Seq and RNA-seq data

**Additional File 7:** Tn-Seq validation

**Additional File 8:** RNA-seq validation

**Additional File 9:** Supplementary Methods: Model assembly and curation; minor metabolite listing; GEO datasets and analysis pipeline.

**Additional File 10:** iSP16 model in SBML format.

**Additional File 11:** Essential genes for T4, 19F, and D39.

